# Allochronic Divergence Driven by Spatial Asynchrony in Precipitation in Neotropical Frogs?

**DOI:** 10.1101/2020.05.05.079210

**Authors:** Carlos E. Guarnizo, Paola Montoya, Ignacio Quintero, Carlos Daniel Cadena

**Affiliations:** Departamento de Ciencias Biológicas, Universidad de los Andes, Bogotá, Colombia; Institut de Biologie de l’École Normale Supérieure (IBENS), ENS, 75005 Paris, France

**Keywords:** Allochrony, genetic distance, nubosity, phenology, rainfall, seasonality, sister species, speciation

## Abstract

The role of geographic barriers in promoting reproductive isolation across space is well understood. Isolation by the time of breeding, however, may also promote population divergence when populations reproduce asynchronically in space, even in the absence of geographic barriers. Few examples exist of divergence due to breeding allochrony, particularly in vertebrates. We tested whether in Neotropical frogs’ asynchrony in precipitation patterns promotes intraspecific genetic divergence, speciation, and regional accumulation of diversity. We assessed the relationship between spatial asynchrony in precipitation and genetic divergence controlling for ecological connectivity across 48 Neotropical frog species. In addition, we examined whether regions within which precipitation regimes are more asynchronous across space have higher species richness and have experienced greater speciation rates. Beyond a generalized expected effect of ecological connectivity on intraspecific genetic divergence, we found that asynchrony in precipitation is positively associated with genetic differentiation in 31% of the species tested, resulting in a significantly positive cross-species effect of asynchrony in precipitation on genetic divergence in a meta-analysis. However, the effect of asynchrony in precipitation on population divergence seems not to scale to macroevolutionary patterns because spatial asynchrony in precipitation was not associated with geographical patterns of species richness or speciation rates. Our results indicate that genetic divergence can be promoted by asynchronous breeding lag in the absence of geographic barriers in species where breeding is associated with water availability, but such effects may not be stable enough through time to influence macroevolutionary patterns.

## Introduction

Understanding the mechanisms underlying spatial variation in biodiversity is a major goal of biogeography and evolutionary biology. The tropics harbor much of the world’s biodiversity, but a thorough understanding of the historical and evolutionary processes underlying such pattern is still lacking. A set of potential drivers focus on mechanisms that enhance tropical rates of speciation, leading to the accumulation of biodiversity over time (Fine, 2015; Fischer, 1960). Whereas the fundamental role that geographical barriers play in promoting speciation by spatially isolating populations has been studied extensively, less attention has been devoted to assessing the role of temporal isolation as a mechanism promoting population divergence (Alexander & Bigelow, 1960; Hendry & Day, 2005; Walter et al., 2017). Namely, if neighbouring populations differ in their reproductive schedules, then gene flow among them is expected to be lower than among populations with overlapping breeding seasons, even in the absence of geographical barriers (Martin, Bonier, Moore, & Tewksbury, 2009; Taylor & Friesen, 2017; Wadgymar & Weis, 2017). Unless the timing of reproductive phenology of a species is entirely plastic, allochronic breeding is expected to promote population differentiation, which could eventually lead to the formation of new species (Taylor & Friesen, 2017). Although there is evidence supporting allochronic speciation across various groups of animals including insects (Santos et al., 2007; Yamamoto & Sota, 2009) and birds (Taylor & Friesen, 2017), the extent and generality of allochronic speciation remains largely unknown, and the role that evolutionary divergence due to asynchronous reproduction may play in establishing spatial patterns of diversity has seldom been considered (Martin et al., 2009).

The “asynchrony of seasons” hypothesis (Martin et al., 2009) posits that spatial asynchrony in breeding seasons may promote genetic divergence among populations, potentially scaling up to influence speciation rates and thereby regional spatial patterns of species richness. Given that species time their reproductive phenology to track annual fluctuations in resource availability, populations from regions varying in the timing of climatic seasonality across space should experience spatial asynchrony in breeding phenology. This, in turn, would result in reduced temporal overlap in reproductive seasons among populations, potentially restricting gene flow and thereby promoting divergence and speciation. Because tropical areas experience much greater spatial asynchrony in environmental factors associated with reproduction (e.g. precipitation and its influence on resource availability) than temperate areas, the asynchrony of seasons hypothesis may partly account for latitudinal differences in species diversity reflecting spatial variation in speciation rates (Martin et al., 2009).

A recent study employing data for several species of birds found support for the asynchrony of seasons hypothesis by assessing whether the degree of precipitation asynchrony among individuals was related to their genetic distance after controlling for ecological connectivity (Quintero, González-Caro, Zalamea, & Cadena, 2014). Assuming that temporal variation in precipitation influences reproductive activity of birds due to its effect on food availability, Quintero et al. (2014) found evidence supporting the hypothesis that spatial asynchrony in breeding seasons influences population differentiation, and thus that current climatic patterns may play a role in driving speciation, contributing to spatial patterns of diversity. However, the authors acknowledged that precipitation may not be the only (nor the most important) driver of breeding schedules of birds and that any effect of precipitation on reproduction would be indirect through its influence on food resources. Given the paucity of data on species’ breeding cycles across different regions, further assessments of the hypothesis that climatic asynchrony drives population divergence are needed, particularly by studying organisms in which climate and reproductive schedules are proximately linked.

Because frogs are expected to closely match their breeding phenology to track temporal variation in precipitation patterns given their strict dependence on water availability for reproduction, they are an ideal group in which to test for an effect of spatial asynchrony in precipitation on population divergence. The hydric physiological restrictions of frogs, which derive mainly from their semipermeable skin and non-amniotic eggs (Lillywhite, 2006), constrain most species to tightly match their reproductive activities with rainy seasons (Kaefer, Montanarin, Da Costa, & Lima, 2012; Saenz, Fitzgerald, Baum, & Conner, 2006; Schalk & Saenz, 2016). Consequently, frogs from populations with asynchronous precipitation regimes should not overlap in their breeding seasons, which may limit gene flow and thereby promote speciation. This effect should be particularly pronounced at lower latitudes because spatial heterogeneity in the timing of climatic seasonality is higher towards the tropics (Duellman, 1995; Martin et al., 2009).

We used DNA sequences and remotely-sensed climatic data to assess whether spatial asynchrony in precipitation seasonality is associated with genetic differentiation in Neotropical frogs. In keeping with previous work, we tested the population-level prediction that precipitation asynchrony should have an effect on genetic distance among individuals of the same species, after controlling for the influence of barriers to dispersal (Quintero et al., 2014). In addition, we tested the macroevolutionary hypothesis that if allochronic speciation driven by climate is recurrent, then lineages inhabiting areas with higher levels of spatial asynchrony in climate should speciate more often, contributing to increased accumulation of species in such areas. All else being equal, this hypothesis predicts that the degree of spatial asynchrony in precipitation within regions should be positively related to regional speciation rates and species richness.

## Methods

### Genetic divergence data

We performed a thorough search of phylogeographic studies of Neotropical frogs and retrieved information on DNA sequences with associated geographic coordinates from GenBank (see Results for the number of species/sequences included). We inspected intraspecific scatterplots of genetic vs geographic distances to make sure there were no highly divergent DNA sequences which may reflect misidentification errors (Guarnizo & Cannatella, 2013). For each species, sequences corresponded to either mitochondrial DNA regions (cytochrome oxidase I, cytochrome b, the control region, NADH dehydrogenase subunit 2) or the 16S ribosomal RNA gene (Table S1). Although markers likely have different substitution rates, this does not affect our intraspecific analyses because each species is analyzed separately. We aligned sequences using the MUSCLE algorithm (Edgar, 2004) and then calculated pairwise genetic distances between all individuals under the best-fit model of nucleotide substitution for each species selected using JModelTest (Posada, 2008) based on the Akaike information criterion (AIC) in the program MEGA 6.0 (Tamura, Stecher, Peterson, Filipski, & Kumar, 2013).

### Asynchrony in precipitation

Asynchrony in precipitation refers to the time lag between peaks in precipitation at two given sites, irrespective of the magnitude of differences in total precipitation between sites. For example, a locality with a precipitation peak occurring in June has greater asynchrony relative to a locality with a precipitation peak in December than a locality where precipitation peaks in April (Fig. 1). Maximum asynchrony occurs between sites with shifted precipitation regimes such that when one site experiences its peak precipitation, the other one experiences the nadir in precipitation (Quintero et al., 2014).

**Figure 1.**
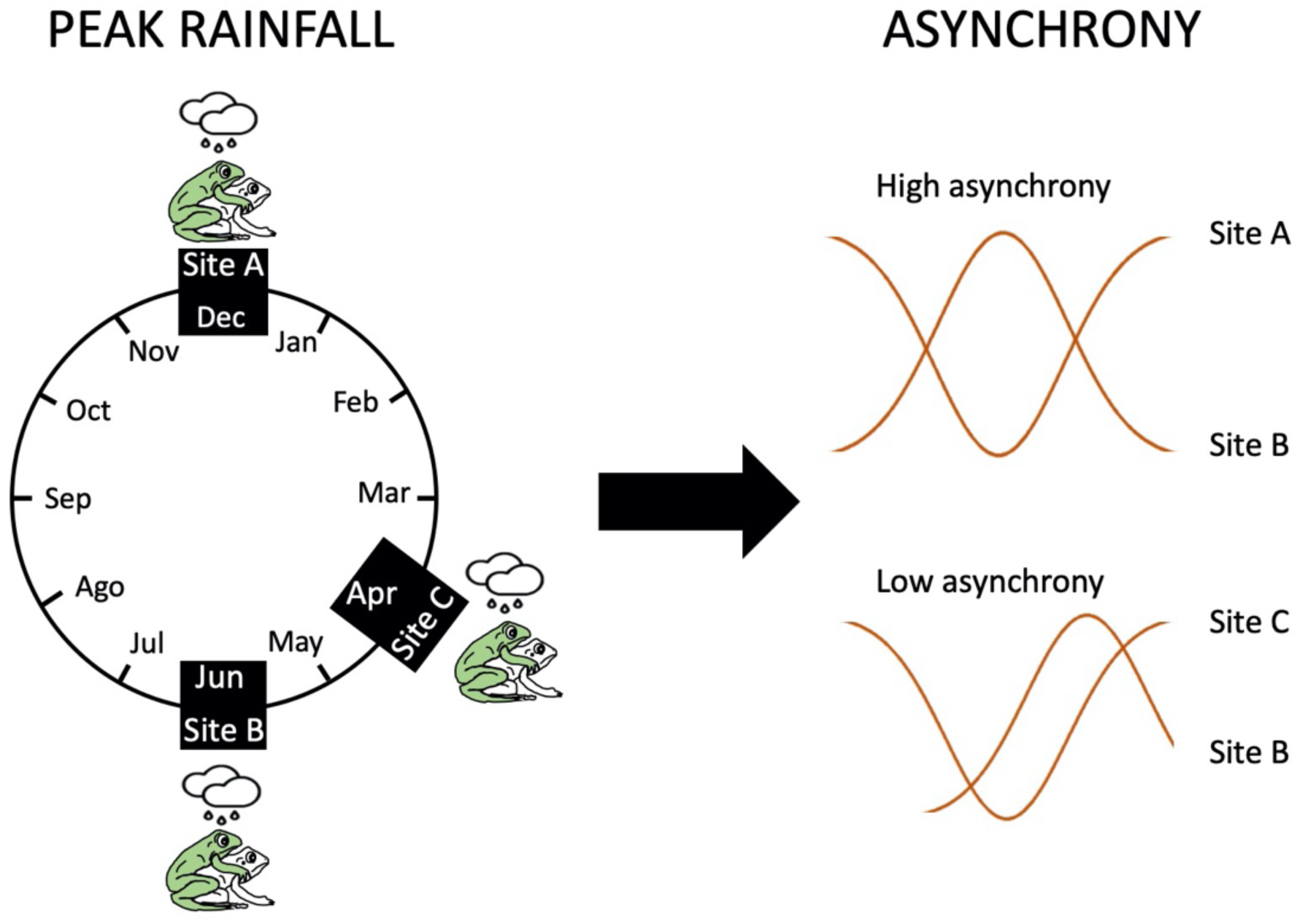
Representation of the asynchrony in precipitation estimation in this study. Left: Frog breeding occurs during peaks in precipitation at three sites: Site A in December, site B in June, and site C in April. Right: Asynchrony in precipitation (represented by sinusoidal curves where peaks correspond to greater precipitation and nadirs to lower precipitation) is larger when comparing sites A and B (December vs. June) and lower between sites C and B (April vs. June).

To estimate asynchrony in precipitation between sites, we followed the approach used by Quintero et al. (2014). We used remotely sensed cloud-cover data to characterize precipitation seasonality based on a public database integrating 15 years of twice-daily remote-sensing cloud observations (MODIS satellite) at 1-km2 resolution (Wilson & Jetz, 2016). Using Fourier transformations, we determined whether each locality experienced a discernible annual (one-peaked) or biannual (two-peaked) rainfall pattern by contrasting the fit of sinusoidal curves with these periodicities to that of a null model. We discarded localities lacking a significant periodic component (i.e. localities with homogeneous precipitation throughout the year). Then, we estimated the lag between the peaks of the sinusoidal curves that best described seasonality among all localities. The maximum lag is six months for annual curves and three months for biannual curves. The estimated lag in peaks was our measure of precipitation asynchrony between localities.

### Accounting for ecological connectivity

We estimated the effect of precipitation asynchrony on genetic divergence while accounting for other factors known to affect population differentiation, namely geographic distance and barriers to dispersal. We used species distribution models to estimate likely dispersal paths given species-specific niche preferences between georeferenced localities, which we then accounted for when assessing the relationship between precipitation asynchrony and genetic differentiation (Quintero et al., 2014). We first obtained geographic coordinates for collecting localities of museum specimens of each species from the Global Biodiversity Information Facility (https://www.gbif.org) and VertNet (http://vertnet.org). We vetted localities by removing coordinates outside range maps defined by experts and the known elevational limits for each species (http://www.iucnredlist.org/). Because range maps constructed based on expert knowledge are reliable at resolutions equal or higher than ca. 1° longitude/latitude (Hurlbert & Jetz, 2007), we incorporated a buffer of 0.5° longitude/latitude around map polygons and discarded georeferenced records located outside. To build species distribution models, we used bioclimatic variables from WorldClim (Hijmans, Cameron, Parra, Jones, & Jarvis, 2005), which describe different aspects of annual temperature and precipitation. Bioclimatic layers that were highly correlated (>70%) were excluded from model estimations using a variance inflation factor analysis in the package ‘USDM’ in R (Naimi, 2015). Subsequently, we estimated a species distribution model for each species using MaxEnt v3.3.3k (Phillips & Dudík, 2008) with 10,000 background points and evaluating model performance with cross-validation using 20% of the data. Finally, we used the inverse of environmental suitability for each species inferred by the models as an ecological resistance matrix from which we estimated the ecological connectivity among localities (i.e. least-cost path distances) using the ‘gdistance’ package for R. We also calculated linear geographic distances (taking into account Earth’s curvature) using the program Geographic Distance Matrix Generator (Ersts, 2011).

### Data analyses at intraspecific level

To assess the effect of precipitation asynchrony, geographic distance, and ecological connectivity on genetic divergence we used multiple-matrix regression with randomization (MMRR), an approach tailored to deal with the non-independence of pairwise distance estimators (Wang, 2013). All distances were standardized (mean = 0, variance = 1) to allow comparison among regression coefficients. We performed MMRR separately on each species using the R script provided by Wang (2013) with 10,000 permutations.

We evaluated an overall, cross-species effect of asynchrony in precipitation on genetic divergence using a phylogenetic meta-analysis (Adams, 2008), where the effect estimated separately for each species was inversely weighted by its associated standard error (i.e. less weight was given to estimates with higher uncertainty) while accounting for the phylogenetic covariance structure. We sampled 100 amphibian phylogenetic trees from the posterior distribution of Jetz and Pyron (2018), a species-complete phylogeny built using sequence data for 15 genes (5 mitochondrial and 10 nuclear) for 4,061 amphibians and with the rest of the species imputed using taxonomic constraints. For the analysis, we pruned all species not considered in our study. For each tree, we estimated this cross-species effect on genetic divergence using Phylogenetic Generalized Least Squares (PGLS) with simultaneous optimization of Pagel’s Lambda under restricted maximum-likelihood using the *gls()* function of the ‘nlme’ package (Pinheiro, Bates, DebRoy, & Sarkar, 2014) for R (Team 2013).

### Species richness and speciation rate

Because the asynchrony of seasons hypothesis posits that seasonal asynchrony in precipitation promotes population differentiation that might scale up to the origin of new species, we expect regions with higher spatial asynchrony in precipitation to have higher speciation rates and, all else being equal, a greater number of species than areas with more spatially uniform precipitation seasonality. We tested these predictions in the Neotropical region (xmin=-118.1333, xmax=-34.79999, ymin=-54.39167, ymax= 33.10834), dividing the region in quadrats of 5 x 5 decimal degrees (grid size of ∼550 x 550 km), a grain size large enough to enable within-quadrat speciation (Kisel & Barraclough, 2010). We estimated the asynchrony in precipitation, species richness, and speciation rates within each one of these quadrats. In the species richness model, we also included productivity and topographic heterogeneity as covariates because these are key factors explaining spatial richness patterns in frogs (Buckley & Jetz, 2007). We describe how we measured these variables below.

We estimated species richness as the sum of all overlapped expert-based maps for anuran species (IUCN 2018) within each 5 x 5 decimal-degree quadrat using ‘raster’ (Hijmans et al., 2015) and ‘maptools’ (Bivand & Lewin-Koh, 2019). R packages. We measured the degree of spatial asynchrony in precipitation within quadrats based on variation in the MODCF seasonality theta parameter (Wilson & Jetz, 2016), which estimates the timing of peak cloudiness in a grid cell as a circular variable ranging from 0 (peak cloudiness on January 1st) to 360 (peak cloudiness on December 31st). Because day of year is a circular variable, we estimated the circular variance of theta values across all the 1 km^2^ cells contained within each 5 x 5 decimal-degree quadrat as our measure of within-region spatial asynchrony in precipitation. The circular variance measures the variation in the angles about the mean direction, ranging from 0 when all data are concentrated at one point (the precipitation peak is at the same time across grid cells within the quadrat) to 1, when the data have broad spread (precipitation peaks are distributed throughout the year within the quadrat). We estimated the circular variance using package ‘circular’ (Agostinelli & Lund, 2017).

To assess frog speciation rates within regions we used the DR metric (Jetz, Thomas, Joy, Hartmann, & Mooers, 2012), which approximates speciation rates under a pure-birth model of diversification and is best interpreted as an estimate of recent speciation rates (Title & Rabosky, 2019). We calculated the DR metric for each species as the median DR across a sample of 100 posterior phylogenetic trees published by Jetz and Pyron (2018). We mapped speciation rates on geography using the harmonic mean value of DR for all species present on each quadrat.

Net primary productivity was obtained from the productivity layer at 0.25 degrees resolution generated by SEDAC (Imhoff et al., 2004) (http://sedac.ciesin.columbia.edu/data/set/hanpp-net-primary-productivity), calculating the mean value in our 5 x 5 decimal degree grid (Fig. S1). This variable is measured in grams of carbon per year, obtained by applying the Carnegie-Ames-Stanford Approach terrestrial carbon model to 17-year average of maximum monthly NDVI (Normalized Difference Vegetation Index) and other climatology data (Imhoff & Bounoua, 2006). Topographic heterogeneity was estimated as the variance in elevation across 1-km^2^ grid cells within each quadrat using the data from STRM (Farr et al., 2007; Fig. S2).

### Data analyses at interspecific level

We ran generalized linear models (GLM) with gaussian-distributed errors to examine the relationship between asynchrony in precipitation and the logarithm of speciation rates, and with Poisson-distributed errors to examine the relationship between asynchrony in precipitation and species richness, using net primary productivity and topographic heterogeneity as covariates. GLMs were performed using the *glm()* function in R. To account for spatial autocorrelation, we used the residual autocovariates (RAC) GLM following Crase et al. (2012), which have shown unbiased coefficients using simulated data. The RAC specifies the relationship between the value of the residuals at a given location and those at neighboring locations. We used the residuals of each GLM model to calculate the RAC as the focal mean value on the first-order neighborhood, and then included it in the model as covariate. The focal mean value was calculated with *focal()* function of ‘raster’ package (Hijmans et al., 2015). We tested the presence of spatial autocorrelation in our data using Moran’s I with the *lm*.*morantest()* function in package ‘spdep’ for all models.

## Results

We evaluated information in ca. 90 papers on Neotropical phylogeography and phylogenetics, and considered 48 species in 16 genera of Neotropical frogs in our study after removing species lacking information on geographic sampling or species occupying landscapes lacking significant periodicity in precipitation patterns (Table S1).

In many species, geographic distance and ecological connectivity were highly correlated (R^2^>0.9). Because MMRR and other multiple regression approaches cannot discriminate individual effects due to collinearity (Wang, 2013), we chose to use ecological connectivity (i.e. least-cost distance) to describe the effect of geography on genetic distances. On average, the full MMRR regression models explained 28% of the variance in intraspecific genetic divergence (Table S2). Ecological connectivity explained most of the variation in genetic distances within species (Fig. S3), with significant effects in 75% of species. Nonetheless, precipitation asynchrony was significantly associated with genetic distances in 31.2% of species (Fig. 2; Table S2). The phylogenetic meta-analysis revealed a significant (albeit weak compared with ecological connectivity) effect of precipitation asynchrony on genetic distances across species (phylogenetic inverse-variance weighted standardized average effect across 100 phylogenetic trees = 0.06 and range = [0.043 - 0.07], average SE = 0.0038, range = [0.0033-0.0079]).

**Figure 2.**
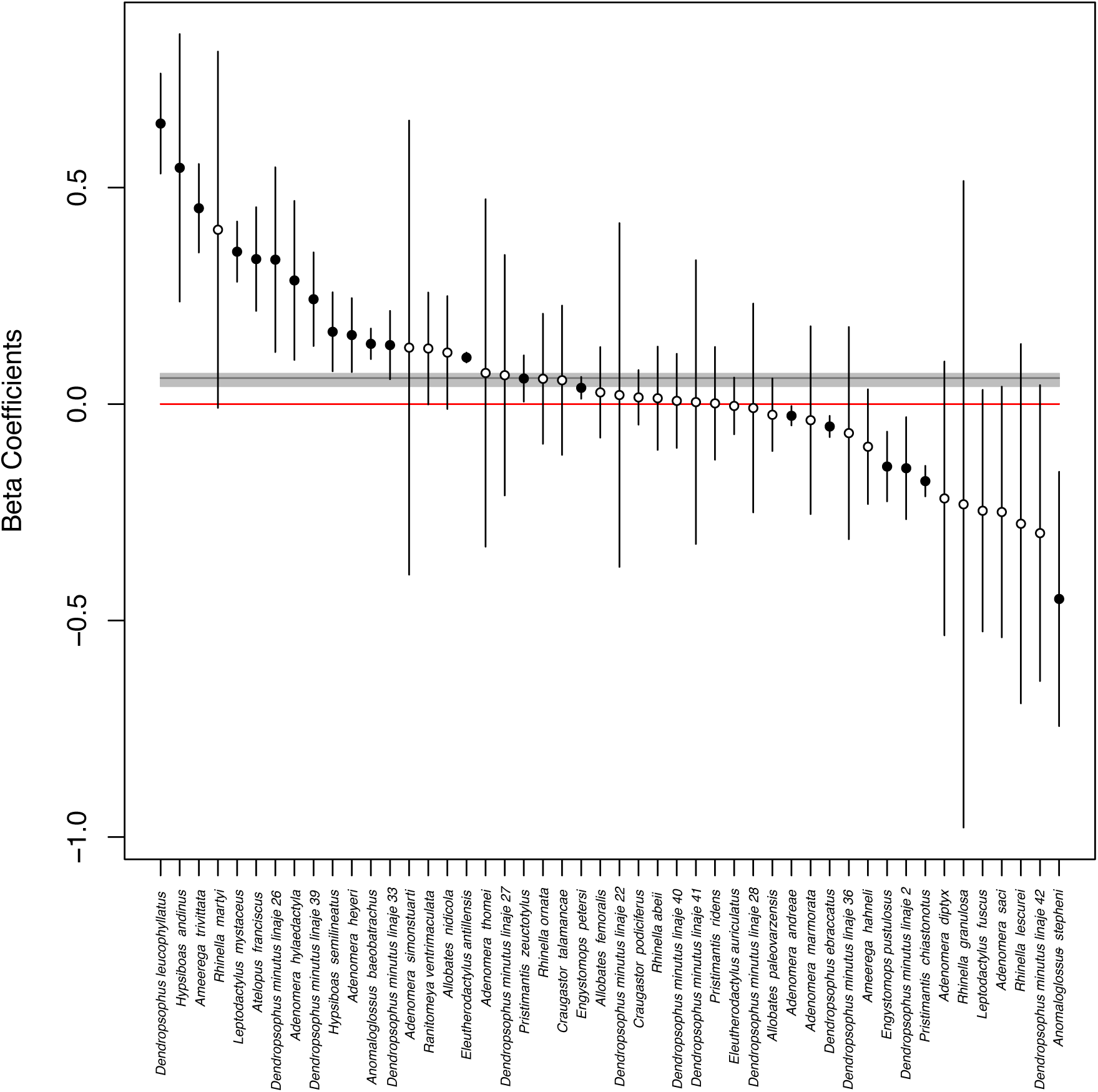
Beta regression coefficients describing the relationship between asynchrony in precipitation and genetic distances, after controlling for the effect of ecological connectivity employing paths of least resistance in 48 species of Neotropical frogs. Black dots indicate a significant association between asynchrony in precipitation and genetic distances (P < 0.05) and white dots indicate non-significant associations between these two variables. Bars indicate 95% confidence intervals. The mean effect of precipitation asynchrony across species using a meta-analysis (horizontal black line) indicates a positive relationship between asynchrony in precipitation and genetic distances. The grey thick horizontal line corresponds to the effect of each of the 100 trees and their confidence intervals (see text). The red reference line corresponds to zero. Because *Dendroposphus minutus* is a species complex, we did not analyse it as a single species, but separated by the highly divergent mtDNA lineages recovered in Gehara et al. (2014), identified with the numbers at the end of each name.

To examine the relationship among precipitation asynchrony, speciation rates and species richness at the regional level, we collected information (geographic and phylogenetic) for 2438 frog species distributed in our study area (Table S3). We found no relationship between spatial asynchrony in precipitation within 5×5 decimal-degree quadrats and mean speciation rates (Fig. 3; glm: slope = 0.005, SE = 0.004, p = 0.146; Moran’s I: -0.112, p = 0.98), but we found a slight negative relationship between spatial asynchrony in precipitation and species richness after accounting for net primary productivity and topographic heterogeneity (Fig. 3; glm: slope = - 0.5420, SE = 0.09827, p-value = 3.48e-08, complete model in Table S4; Moran’s I: -0.08, p = 0.95). In this latter model, the covariates (primary productivity and topographic heterogeneity) also show a significant positive, although slight, relationship with species richness (Table S4).

**Figure 3.**
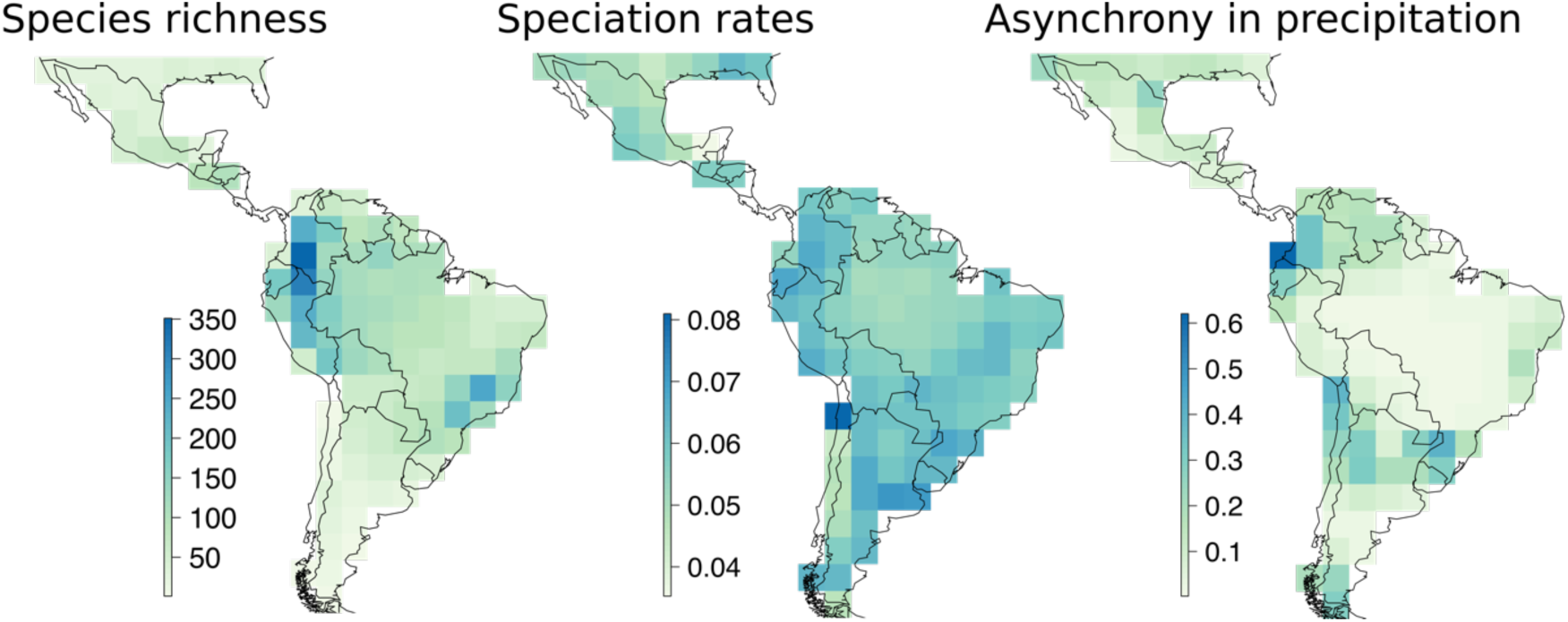
Maps depicting frog species richness, speciation rate, and spatial asynchrony in precipitation within 5×5 decimal degree quadrats. There is no spatial relationship between the asynchrony in precipitation and species richness or speciation rate after controlling for the positive effects of topographic heterogeneity and net primary productivity.

## Discussion

As predicted by the asynchrony of seasons hypothesis (Martin et al., 2009), we found that greater asynchrony in precipitation is associated with higher genetic differentiation among populations of Neotropical frogs after controlling for effects of geography/ecology and phylogeny. We found this pattern in 16 of the species tested, which led to a significant –albeit weak– cross-species effect in a phylogenetic meta-analysis considering data for 48 species. Contrary to predictions of the hypothesis, however, frogs in geographic regions in which precipitation regimes are more variable in space do not appear to have experienced higher speciation rates nor accumulated higher richness than in regions where precipitation is more spatially synchronic. Mismatches in the breeding phenology of populations linked to spatial patterns of asynchrony in precipitation have been scarcely explored as a potential driver of diversification (Martin et al., 2009; Quintero et al., 2014). In particular, aside from an earlier study on New World birds showing significant associations between asynchrony in precipitation and genetic distances (Quintero et al., 2014), we are unaware of comprehensive tests of the asynchrony of seasons hypothesis (Martin et al., 2009) and no multispecies analysis has tested its predictions in organisms directly relying on water availability to reproduce.

Although aspects of breeding biology are largely unknown for many Neotropical species, seasonal variation in precipitation is known to correlate with reproductive activity in frogs, including some of the species in this study (Donnelly & Guyer, 1994; Kaefer et al., 2012; Moreira & Lima, 1991). Assuming that precipitation is a proximate surrogate for breeding activity in frogs, our study expands the understanding of the importance of reproductive timing as a factor promoting lineage divergence at two different evolutionary scales: intraspecific genetic divergence, and speciation rates. The positive association between spatial asynchrony in precipitation and genetic distances in several of the species tested is consistent with the hypothesis that spatial variation in water availability may cause spatial asynchrony in breeding activity, which may in turn result in reduced gene flow among populations from regions with asynchronous precipitation regimes. However, our results suggest that this mechanism of population divergence does not seem to scale up in driving rates of species origination and thus does not enhance the accumulation of regional diversity.

A previous study in a frog species found evidence consistent with the hypothesis that differences in timing of reproduction mediated by water availability drive intraspecific genetic differentiation. Breeding activity of *Rana arvalis* in Europe and Asia is linked to ice melt, which occurs at different times in different populations, resulting in spatial mismatches in the availability of ponds and hence in breeding phenology (Richter-Boix, Quintela, Kierczak, Franch, & Laurila, 2013). Differences in breeding times are positively associated with genetic distances in *R. arvalis*, suggesting that populations may diverge owing to allochrony in the absence of geographic barriers to gene flow (Richter-Boix et al., 2013). Our study found a similar effect across species in association with spatial variation in precipitation in the Neotropics, suggesting that models describing population genetic differentiation in amphibians (Zeisset & Beebee, 2008) would benefit by considering mechanisms beyond geographic distance and geographic barriers.

For the asynchrony of seasons hypothesis to hold, the driest season at any given locality should experience less precipitation than the minimum required for breeding. If two regions are highly asynchronous in their rainfall patterns yet experience precipitation magnitudes above the minimum threshold necessary for breeding, then genetic divergence between regions should not be expected. Another assumption of the hypothesis is that breeding phenology is not entirely caused by the environment; if dispersing individuals may readily accommodate their breeding condition to adjust to local availability of food or water through phenotypic plasticity, then reduced gene flow among climatically asynchronous populations is not expected (reviewed by Quintero et al. 2014). Breeding times in salmonid fish and some birds are strongly heritable (Price, Kirkpatrick, & Arnold, 1988), such that species breed at the same time of year regardless of environmental fluctuations. In amphibians, the heritability of breeding times is unclear and may vary across taxa. For example, the onset of breeding in salamanders is determined primarily by environmental factors partially constrained by underlying life-history trade-offs (Semlitsch, Scott, Pechmann, & Gibbons, 1993). In contrast, traits such as spawning time in the frog *Rana temporaria* reflect local adaptation, implying a heritable component to reproductive phenology (Phillimore, Hadfield, Jones, & Smithers, 2010). Because we have no information about the heritability of breeding times (or phenological cues) in our species, it would be important to evaluate it in future studies to further test the asynchrony of seasons hypothesis.

Frogs exhibit a diversity of reproductive modes depending on the availability of water and competition for resources (Haddad & Prado, 2005), with two major breeding categories at the extremes of a continuum: explosive and prolonged (Wells, 1977). Explosive breeders are usually found in dry habitats where water is scarce during most of the year; after heavy rains, individuals congregate and mate in recently formed water bodies during one or few nights. On the contrary, prolonged breeders mate throughout the year and usually inhabit sites maintaining humidity through time. Even though some species of frogs are more inclined to be explosive or prolonged, many appear to shift between reproductive modes depending on environmental factors (Touchon & Warkentin, 2008; Wells & Schwartz, 2006). Intuitively, one would expect to find stronger effects of precipitation asynchrony on genetic differentiation in species closer to the explosive end of the continuum because their breeding is temporally restricted, and therefore, temporal isolation among regions is more probable. If, however, rainy periods are unpredictable, then frog species may be selected for plasticity and reproduce opportunistically whenever it starts raining, as observed in birds inhabiting unpredictable environments (Hau, Wikelski, Gwinner, & Gwinner, 2004). Also, one would expect that frogs laying eggs on microhabitats requiring more water should show stronger effects of precipitation asynchrony on genetic differentiation. For example, species laying eggs on vegetation need moisture for clutch development and high water levels for larvae to drop safely into ponds or streams (Gottsberger & Gruber, 2004), whereas direct-development species lay eggs in leaf litter where they are more buffered from fluctuations in moisture driven by precipitation. Despite the expectation that reproductive mode and microhabitat would mediate in the association between precipitation and breeding time (hence genetic divergence), species in which we encountered a positive and significant relationship between asynchrony in precipitation and genetic distances belonged to a variety of frog families characterized by very different reproductive biology (Bufonidae, Leptodactylidae, Craugastoridae Dendrobatidae, Hylidae). For instance, among the ten species showing the strongest association between asynchrony in precipitation and genetic distances were frogs with direct development (*Pristimantis zeuctotylus*), aquatic eggs (*Rhinella martyi, Dendropsophus minutus*), semiterrestrial eggs deposited on leaves (*Dendropsophus leucophyllatus*), eggs on water tanks on epiphytic plants (*Ranitomeya ventrimaculata*), and foam nests built on the ground (*Leptodactylus mystaceus, Adenomera hylaedactyla*). Therefore, populations of species with particular reproductive modes do not appear more likely to diverge owing to spatial asynchrony in precipitation.

Our multiple regression models indicated that the effect of ecological connectivity on within-species genetic distances, was on average, three times larger than the effect of precipitation asynchrony. This result confirms the substantial role that spatial isolation (particularly in topographically heterogeneous regions) has on the genetic differentiation of Neotropical amphibians. Because topography and precipitation patterns are closely linked (Viale et al., 2019), separating the independent effects of these variables on population differentiation may be difficult. For example, the eastern slopes of cordilleras in the Northern Andes are in general more humid than the western slopes due to trade winds and rainshadow effects (with some exceptions such as the Chocó region). If a species exists on both sides of the Andes, then distinguishing the effects of mountains, precipitation and their interaction as leading sources of genetic divergence is challenging. We believe that in most cases topography is more important than precipitation in driving diversification owing to physical and physiological restrictions to dispersal across geographic barriers (Janzen, 1967). Nonetheless, our results indicate that asynchrony in precipitation may be causally linked to divergence of populations in some species of frogs, suggesting that future work should consider spatial allochrony mediated by climate as a likely driver of divergence.

A salient conclusion of our work is that support for the asynchrony of seasons hypothesis seems to be observable only at a microevolutionary scale because asynchrony in precipitation was associated positively with genetic distances, but not with speciation rates nor species richness. This may reflect temporal instability of spatial patterns in precipitation driven by factors such as orbital precession (Merlis, Schneider, Bordoni, & Eisenman, 2013), climatic change associated with glacial cycles (Whitney et al. 2011), or climate modulation caused by the uplift of the Andes (Insel, Poulsen, & Ehlers, 2010). Such historical events likely modified geographic patterns in precipitation asynchrony, which may have been stable enough to allow for intraspecific genetic differentiation in Neotropical frogs, but not to complete speciation across asynchronous regions in the absence of physical barriers to gene flow. A similar explanation involving climatic instability has been advanced to account for the lack of evidence of parapatric speciation across habitat ecotones despite numerous cases of adaptive divergence of populations in such ecological settings (T. Smith, R. Wayne, D. Girman, 2005). Unfortunately, historical data on temporal variation in precipitation regimes across regions which allow us to consider such dynamics in our study are unavailable.

Historically, topography and temperature gradients have been the main variables used to study patterns of genetic variation and diversification of frogs. Our results suggest that genetic divergence can be promoted in the absence of geographic barriers or temperature gradients in species where breeding is associated to water availability. As information on current and past precipitation patterns becomes more accessible through remote sensing and public databases, explorations of the phenology and heritability of breeding times of species will become more important to provide a fine grain understanding of how precipitation can promote genetic divergence, and perhaps, Neotropical diversification.

## Acknowledgments

The authors would like to thank Melissa Hernández and Gabrielle Genty for invaluable help compiling DNA sequences and locality data from the literature, and Laura Céspedes for helping with the SDM’s. We also thank Elkin Tenorio, Sebastián González and the Evolvert and Biomics labs at Universidad de los Andes for helpful comments and suggestions that greatly improved this manuscript.

