## Supplementary material for "Allochronic Divergence Driven by Spatial Asynchrony in Precipitation in Neotropical Frogs?": Suplemental figures

**Figure S1.** Net primary productivity map at 0.25° resolution generated by SEDAC in grams of carbon (Imhoff et al. 2004). See text for details.

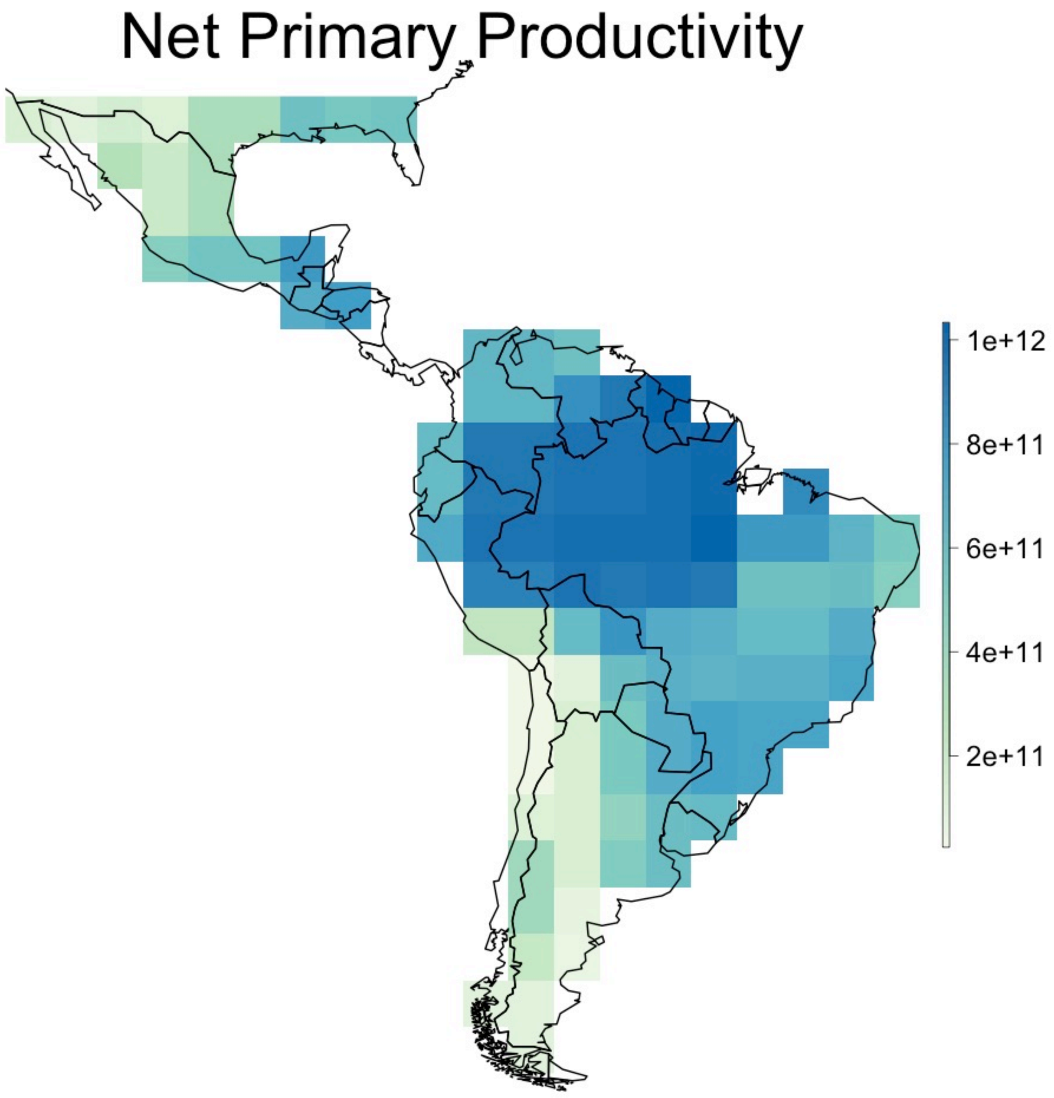

**Figure S2.** Topographic heterogeneity map estimated as the variance in elevation across 1-km2 grid cells. Colors towards blue indicate more heterogeneity. See text for details.

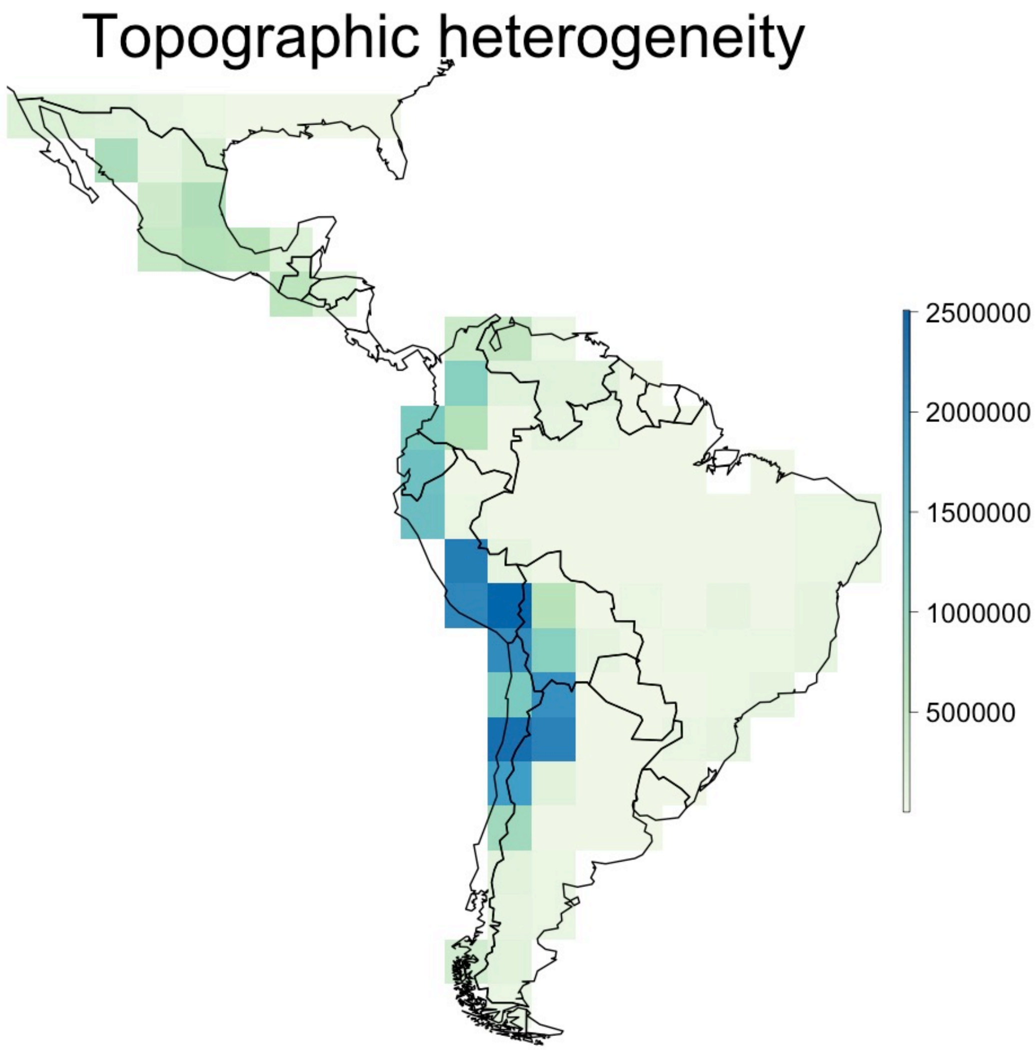

**Figure S3.** Beta coefficients of the multiple regression (MMRR) between ecological connectivity (Least Cost Path distances) and genetic distances within species, after controlling for the effect of precipitation asynchrony. Black dots indicate a significant association between asynchrony in precipitation and genetic distances (P < 0.05) and white dots indicate non-significant associations between these two variables. Bars indicate 95% confidence intervals. The horizontal red line corresponds to zero.

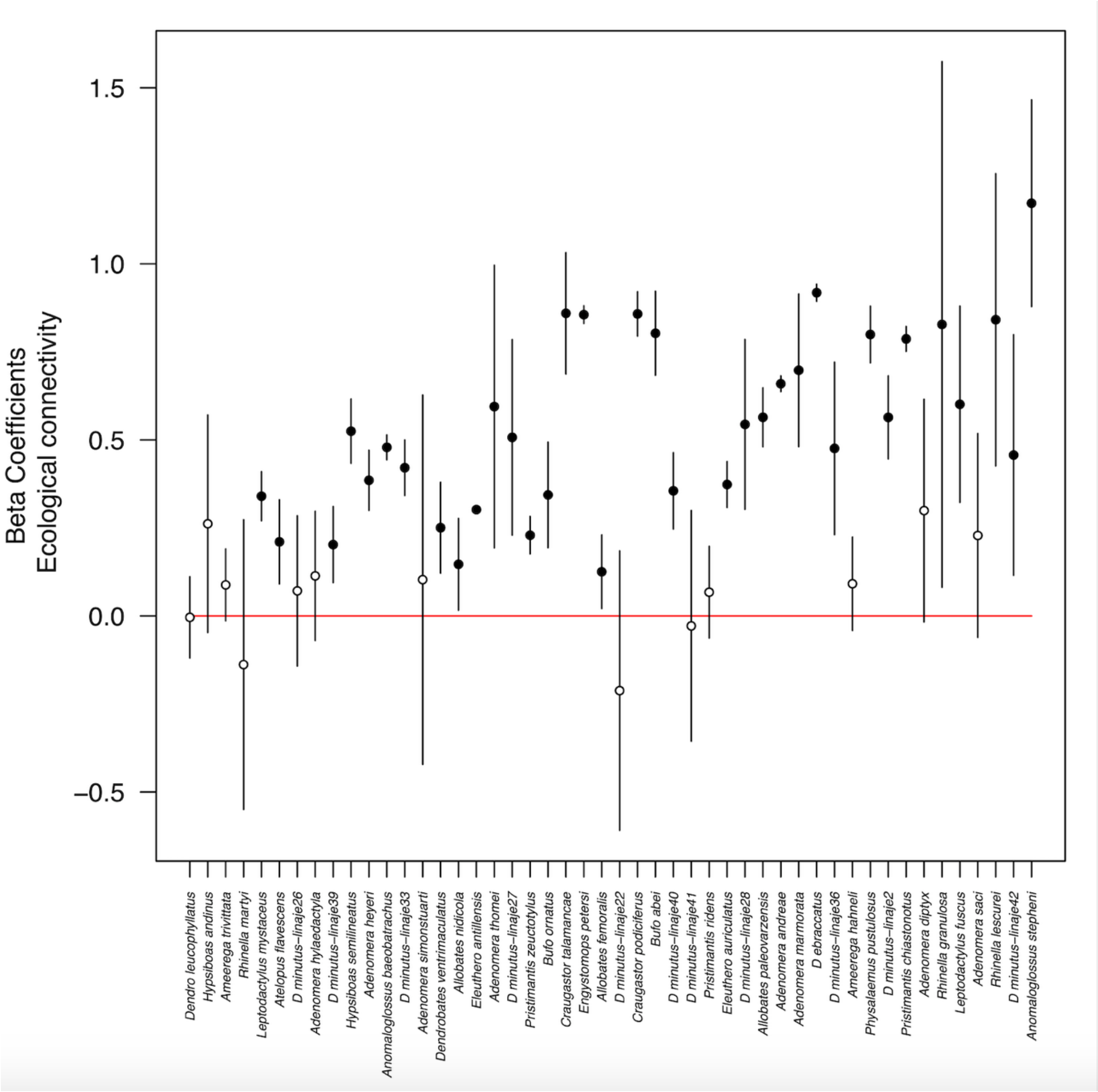
